# Aromatic acid metabolism in *Methylobacterium extorquens* reveals interplay between methylotrophic and heterotrophic pathways

**DOI:** 10.1101/2025.03.22.644763

**Authors:** Alekhya M. Govindaraju, Norma Cecilia Martinez-Gomez

**Affiliations:** Department of Plant and Microbial Biology, University of California, Berkeley, Berkeley, CA, USA

## Abstract

Efforts towards microbial conversion of lignin to value-added products face many challenges because lignin’s methoxylated aromatic monomers release toxic C_1_ byproducts such as formaldehyde. The ability to grow on methoxylated aromatic acids (e.g., vanillic acid) has recently been identified in certain clades of methylotrophs, bacteria characterized by their unique ability to tolerate and metabolize high concentrations of formaldehyde. Here, we use a phyllosphere methylotroph isolate, *Methylobacterium extorquens* SLI 505, as a model to identify the fate of formaldehyde during methylotrophic growth on vanillic acids. *M. extorquens* SLI 505 displays concentration-dependent growth phenotypes on vanillic acid without concomitant formaldehyde accumulation. We conclude that *M. extorquens* SLI 505 overcomes potential metabolic bottlenecks from simultaneous assimilation of multicarbon and C_1_ intermediates by allocating formaldehyde towards dissimilation and assimilating the ring carbons of vanillic acid heterotrophically. We correlate this strategy with maximization of bioenergetic yields and demonstrate that formaldehyde dissimilation for energy generation rather than formaldehyde detoxification is advantageous for growth on aromatic acids. *M. extorquens* SLI 505 also exhibits catabolite repression during growth on methanol and low concentrations of vanillic acid, but no diauxie during growth on methanol and high concentrations of vanillic acid. Results from this study outline metabolic strategies employed by *M. extorquens* SLI 505 for growth on a complex single substrate that generates both C_1_ and multicarbon intermediates and emphasizes the robustness of *M. extorquens* for biotechnological applications for lignin valorization.

## Introduction

Lignin, a major component of woody plant cell walls, is one of Earth’s most abundant renewable carbon sources. It comprises a complex network of polycyclic aromatic polymers that provide rigidity for growing plants and acts as a barrier against harsh weather and grazing herbivores^1,2^. Microbial degradation of lignin is interesting both from ecological and biotechnological perspectives, as increased understanding of these processes relate to efficient carbon cycling in natural ecosystems as well as the exploitation of lignin as a feedstock for petrochemical production^2,3^.

Microbial degradation of lignin-derived methoxylated aromatic acids (e.g., vanillic acid, ferulic acid, protocatechuic acid) is distributed across soil and plant microorganisms^2,4,5^. While variations of aromatic acid degradation modules exist^6^, vanillic acid is commonly used as a model for investigating aromatic acid degradation^7–9^. Aerobic growth on vanillic acid proceeds through the following enzymatic reactions: ferulic acid is oxidized to vanillic acid (*ech*), which is demethylated to protocatechuic acid (*vanAB*) and produces formaldehyde as an obligate byproduct. Protocatechuic acid serves as a substrate for the β-ketoadipate pathway. It undergoes a series of ring cleavage steps (*pcaHG, pcaB, pcaC, pcaD*) to generate β-ketoadipate, which is converted to succinyl-CoA and acetyl-CoA (*pcaIJ, pcaF*), common building block metabolites for the TCA cycle and other assimilatory cycles (**Figure 1A**, pathway indicated in blue)^10,11^.

**Figure 1.**
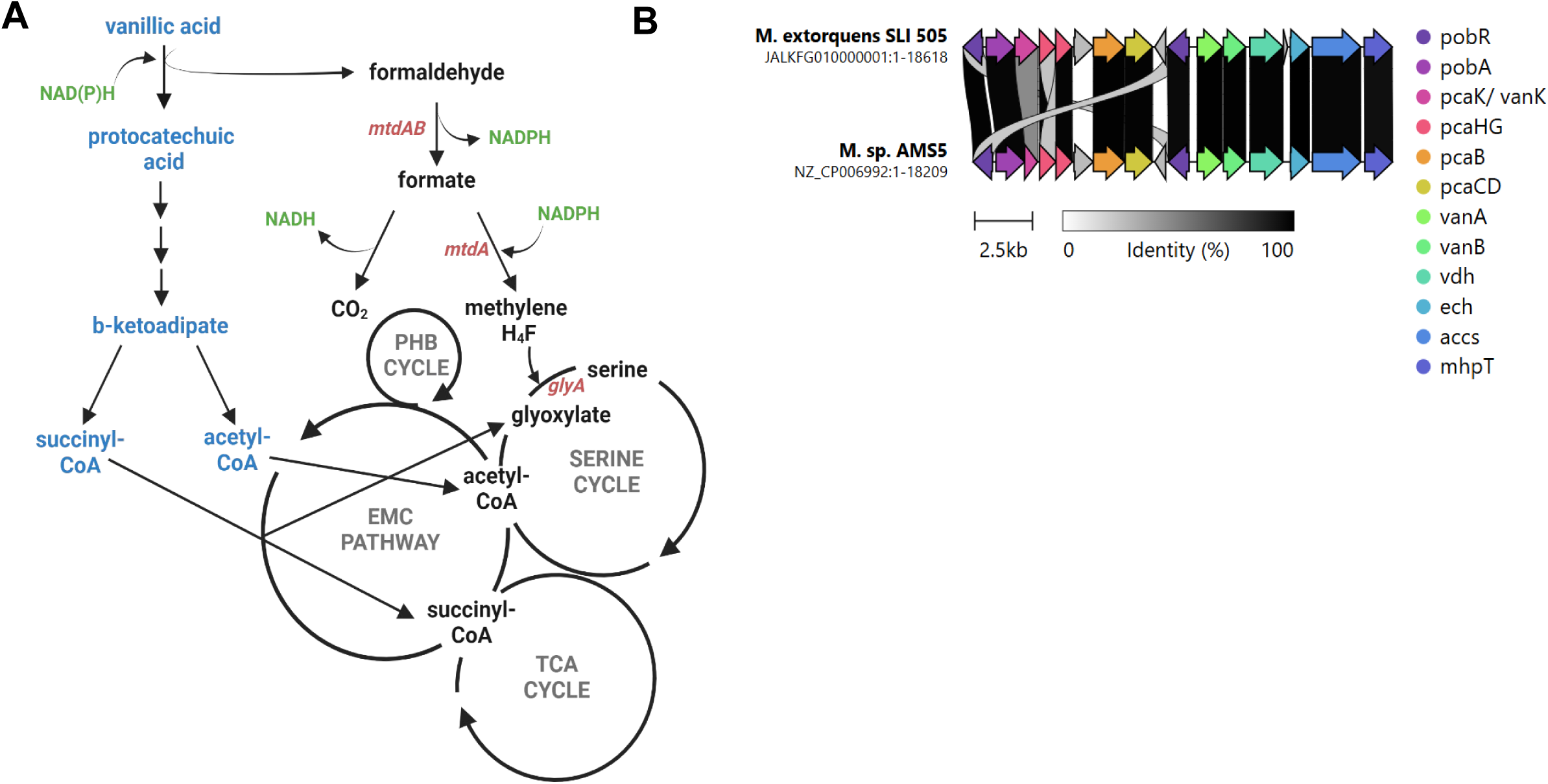
Annotations of aromatic acid gene island in *Methylobacterium extorquens* predicts how aromatic acid metabolism integrates into facultative methylotrophy. **A**. Predicted metabolic map of growth on aromatic acid in *M. extorquens* SLI 505. Aromatic acid pathways shown in blue, relevant genes in red. **B**. Aromatic acid gene island in *M.extorquens* SLI 505 and *Methylobacterium sp.* AMS5. Comparative analysis generated using CAGECAT cluster analysis toolbox

The obligate production of formaldehyde in the conversion of vanillic acid to protocatechuic acid has been hypothesized to be a major bottleneck in the efficient degradation of methoxylated aromatic acids^12–15^. Formaldehyde is a highly toxic electrophile that can crosslink with proteins and DNA, inhibiting cell growth^16^. Previous studies have shown that *Pseudomonas putida* KT2440^17,18^ and *Rhodococcus jostii* RHA1^19^, both model organisms for aromatic acid metabolism, release appreciable amounts of formaldehyde during growth on vanillic acid. Formaldehyde excretion correlates with cell death in these organisms despite both encoding native formaldehyde detoxifying mechanisms^18^. To overcome the rate-limiting step of C_1_ byproduct toxicity, there have been efforts to engineer microbes such as *Escherichia coli*, *Corynebacterium glutamicum*, and *Burkholderia cepacia* for enhanced aromatic acid degradation through substrate channeling, compartmentalization, and introduction of novel formaldehyde oxidation pathways^5,12^.

The majority of microorganisms are highly susceptible to formaldehyde toxicity, but methylotrophic bacteria have evolved robust formaldehyde detoxification mechanisms. Methylotrophs are defined by their ability to utilize reduced carbon compounds lacking carbon-carbon bonds (e.g., methane, methanol, methylamine) as their sole source of carbon and energy^20,21^. Recently, the ability to degrade methoxylated aromatic acids has been identified in specific clades of methylotrophic bacteria. Still, the extent to which methoxydotrophy–the metabolism of methoxy groups of aromatic compounds– occurs is not widely understood^19^. During methylotrophic growth in the model, pink-pigmented facultative methylotroph, *Methylobacterium extorquens*, coupled redox reactions in the periplasm oxidize primary C_1_ substrates (methanol, methylamine, etc) to formaldehyde and transfer the resulting electrons to cytochromes directly linked to the electron transport chain for energy conservation^20,22^. Formaldehyde is transported from the periplasm to the cytoplasm, where it is immediately covalently attached to a tetrahydromethanopterin (H_4_MPT) carbon carrier and oxidized to formate through a series of steps that generate NAD(P)H as reducing power^20,22^. Formate serves as a branchpoint between assimilation and dissimilation. Formate can be oxidized to CO_2_ with the concomitant generation of NADH via formate dehydrogenases or assimilated via a partial reduction facilitated by a tetrahydrofolate carbon carrier to enter the serine cycle for assimilation (**Figure 1A**, pathway in black)^23^. Methylotrophic growth is considered limited by NADPH, as the primary source of NADPH necessary to assimilate formate into the serine cycle is the oxidation of formaldehyde to formate in the preceding steps^20,23,24^.

Degradation of methoxylated aromatic acids by methylotrophs yields both C_1_ (formaldehyde) and multicarbon (acetyl-CoA, succinyl-CoA) intermediates – an example of a single complex substrate that produces two distinct intermediates that can be assimilated into metabolic pathways via different entry points (**Figure 1A**). Many methylotrophs, including members of the *extorquens* clade, are facultative organisms capable of utilizing multicarbon compounds as a carbon and energy source by feeding intermediates directly into the TCA cycle for assimilation^20,21^. Because the serine cycle, used for the assimilation of C_1_ intermediates, and the TCA cycle, used for the assimilation of multicarbon intermediates, share enzymatic reactions but run in opposite directions, simultaneous assimilation of C_1_ and multicarbon compounds cannot occur^21,24^. Previous studies have demonstrated that when methylotrophs are grown on succinate (multicarbon) and methanol (C_1_) simultaneously, they allocate the C_1_ substrate to dissimilation and the multicarbon substrate to assimilation^25^ – a strategy that overcomes diauxic growth while maximizing bioenergetic yields during co-substrate growth.

In this study, *Methylobacterium extorquens* SLI 505, a recent isolate from the soybean phyllosphere^26^, is used as a model to understand how methylotrophs overcome bottlenecks surrounding methylotrophic and heterotrophic pathway operation in their metabolism of aromatic acids. *M. extorquens* SLI 505 upregulates its methylotrophic and heterotrophic machinery in response to increasing concentrations of vanillic acid, yet formaldehyde is not assimilated during growth. Further, formaldehyde detoxification only enables optimization of carbon utilization at the level of growth rate and not yield, despite that formaldehyde detoxification is efficient even during growth on high concentrations of vanillic acid. Finally, catabolite repression is observed when methanol and low concentrations of vanillic acid are available. Our data suggests that currency metabolites such as NAD(P)H may play a role in defining the distribution of carbon between assimilation and dissimilation.

## Results

### Methylobacterium extorquens SLI 505 is an optimal model for investigating native vanillic acid metabolism in methylotrophic bacteria

Previously, our lab isolated 78 *Methylobacterium extorquens* strains capable of robust growth on vanillic acid from the phyllosphere of soybeans, collectively referred to as SLI (**<u>S</u>**oybean **<u>L</u>**eaf **<u>I</u>**solate) strains^26^. Whole-genome sequencing through the Joint Genome Institute of all SLI strains capable of growth on aromatic acids revealed the presence of a horizontally-transferred ∼19-kb cluster that encodes enzymes necessary for vanillate (VanAB), ferulate (Ech), and protocatechuate (PcaHG, PcaB, PcaC, PcaD, PcaIJ, PcaF) metabolism via the β-ketoadipate pathways described above. This cluster is hereafter referred to as the aromatic gene island. Of the SLI strains capable of growth on vanillic acid, *M. extorquens* strain SLI 505 had the most reproducible growth phenotypes in preliminary experiments and was thus selected as the model organism by which to evaluate vanillic acid metabolism in the *M. extorquens* SLI community.

The CAGECAT toolbox^27^ was used for comparative analysis of aromatic acid gene islands in *M. extorquens* SLI 505 and *Methylobacterium sp*. AMS5, the only other reported member of the *extorquens* clade of *Methylobacterium* capable of growth on aromatic acids (**Figure 1B**)^19^. SLI strains grow faster, to a higher final OD_600_, and more reproducibly than *M. sp.* AMS5 on vanillic acid^26^. Despite differences in growth phenotypes, the aromatic acid gene islands of both strains share high sequence identity and organization. In contrast, homologous genes for aromatic acid degradation for *Methylobacterium* members of the *aquaticum* and *nodulans* clades^19^– in which this metabolic capacity is more widespread– as well as in *Pseudomonas putida* KT2440^28^ – a non-methylotrophic bacteria with well-characterized aromatic acid metabolism– exist as disparate clusters across the genome and lack the consolidated organization exhibited within the *extorquens* clade.

### Methylotrophs exhibit growth defects at high concentrations of vanillic acid without concomitant formaldehyde accumulation

To establish how *M. extorquens* SLI 505 grows on a range of vanillic acid concentrations, this strain was grown in minimal media with increasing concentrations (0.25 to 20 mM) of vanillic acid supplemented as the sole carbon and energy source. *M. extorquens* SLI 505 exhibits a concentration-dependent growth phenotype on vanillic acid. At substrate concentrations from 0.5 mM to 6 mM vanillic acid, growth rates and maximum OD_600_ values increase in a pattern that correlates with the amount of carbon provided. Growth rates decrease at substrate concentrations higher than 6 mM vanillic acid (**Figure 2B**), and maximum OD_600_ values stabilize at approximately 10-12 mM vanillic acid; despite that growth is reported at higher substrate concentrations, maximum OD_600_ values do not increase (**Figure 2C**), and growth rates decrease (**Figure 2B**). Based on the growth phenotypes for *M. extorquens* SLI 505 in **Figure 2**, the substrate threshold for vanillic acid was subsequently classified as “low concentration” (0.5 - 6 mM), “medium concentration” (7 - 9 mM), and “high concentration” (10 - 15 mM) for ease of reference throughout this study.

**Figure 2.**
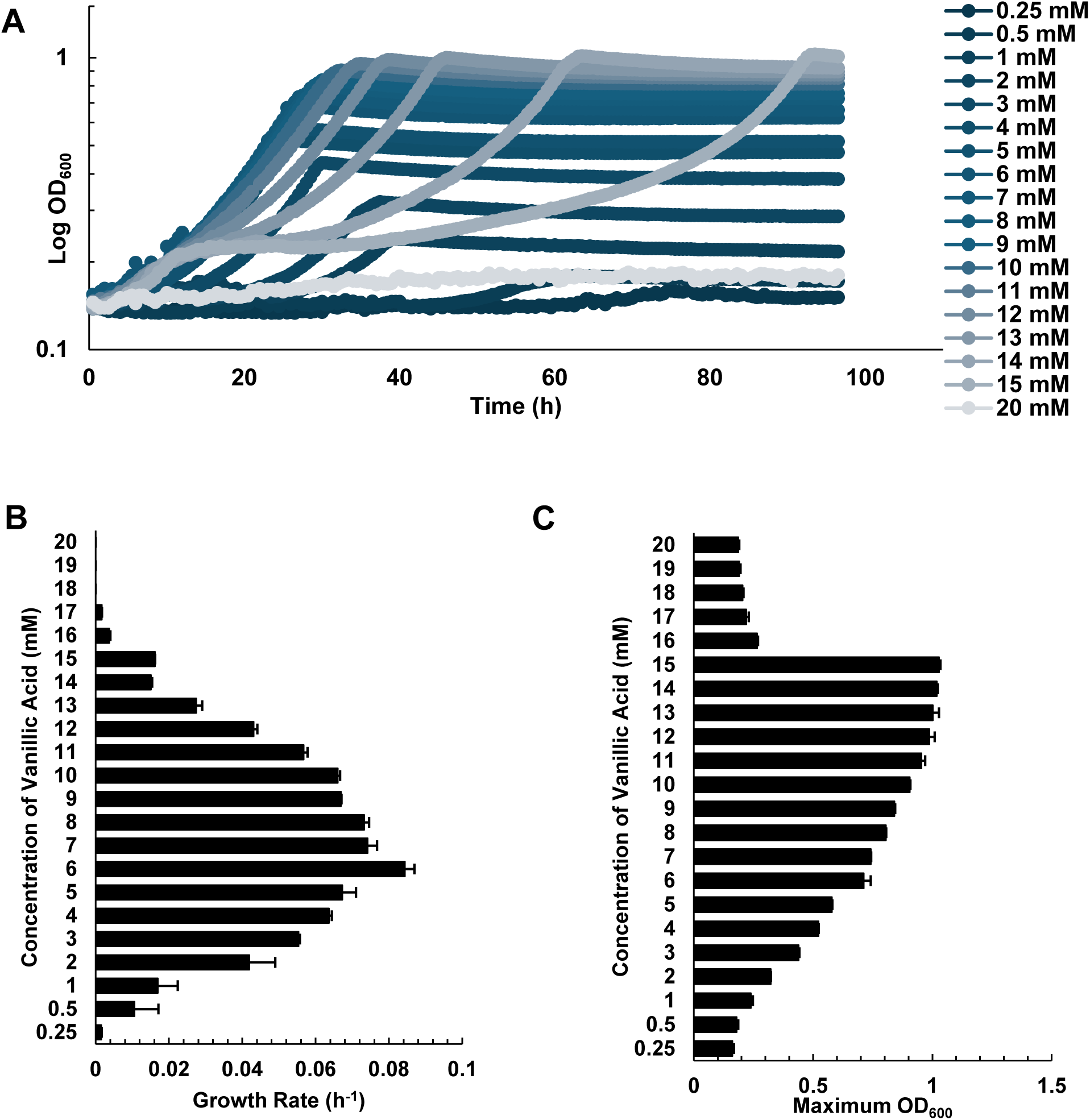
Growth phenotypes of *Methylobacterium extorquens* SLI 505 on minimal media supplemented with 0-20 mM vanillic acid. **A.** Growth curves of *M. extorquens* SLI 505 on 0.25-20 mM vanillic acid. Each curve represents an average of two replicates. **B.** Growth rates of *M. extorquens* SLI 505 on 0.25-20 mM vanillic acid. Error bars indicate standard deviation of two replicates. **C.** Maximum OD_600_ of *M. extorquens* SLI 505 on 0.25-20 mM vanillic acid. Error bars indicate standard deviation of two replicates.

It was hypothesized that the decrease in growth at higher concentrations of vanillic acid was due to formaldehyde accumulation, as shown for non-methylotrophs which succumb to formaldehyde toxicity at lower substrate concentrations^18,29^. To assess this, intracellular formaldehyde concentrations in a range of vanillic acid concentrations were measured (**Table 1**). Appreciable concentrations of formaldehyde^30^ were not detected during growth on 5-12 mM vanillic acid. The lack of formaldehyde accumulation at any vanillic acid concentration is in sharp contrast to the formaldehyde accumulation observed in non-methylotrophs, even at lower vanillic acid concentrations, and emphasizes the robustness of methylotrophs for mitigating formaldehyde toxicity during growth on aromatic acids.

**Table 1.**
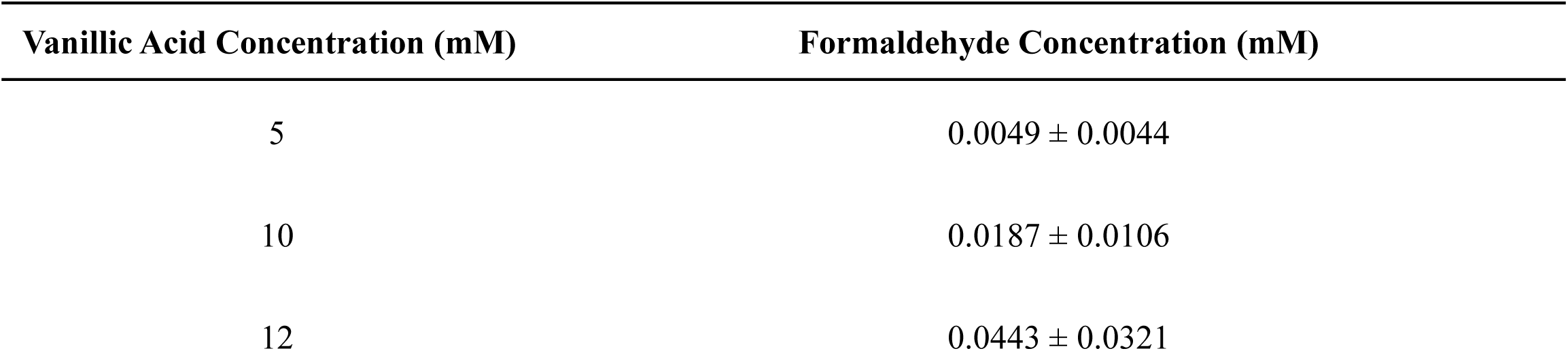
Intracellular formaldehyde concentrations of *M. extorquens* SLI 505 during growth on vanillic acid.

| Vanillic Acid Concentration (mM) | Formaldehyde Concentration (mM) |
| --- | --- |
| 5 | $0.0049 \pm 0.0044$ |
| 10 | $0.0187 \pm 0.0106$ |
| 12 | $0.0443 \pm 0.0321$ |

### Methylotrophs do not assimilate formaldehyde during growth on vanillic acid

If formaldehyde is not accumulating during growth of methylotrophs of vanillic acid, it must be assimilated, dissimilated, or excreted into the extracellular medium. We know that the latter is not occurring based on measurements of extracellular formaldehyde at the indicated substrate concentrations in **Table 1** that found negligible or zero amounts of formaldehyde (data not shown), leaving two other possible fates. To track formaldehyde assimilation into biomass or dissimilation towards CO_2_, vanillic acid with a ^13^C label on the methoxy carbon, hereafter referred to as ^13^C-vanillic acid (**Figure 3A**), was used for formaldehyde labeling studies. Through ^13^C fingerprinting, the incorporation of ^13^C carbon into proteinogenic amino acids was measured by liquid chromatography-mass spectrometry and used as a proxy for determining if carbon from formaldehyde was assimilated towards biomass. **Figure 3B** reports the mass isotopomer distribution in measurable amino acids, where the M-number indicates the number of ^13^C carbons identified in each amino acid. ^13^C carbon was not detected in most amino acids in substantial amounts (**Figure 3B)**, indicating that formaldehyde generated during growth on aromatic acids is not assimilated towards biomass.

**Figure 3.**
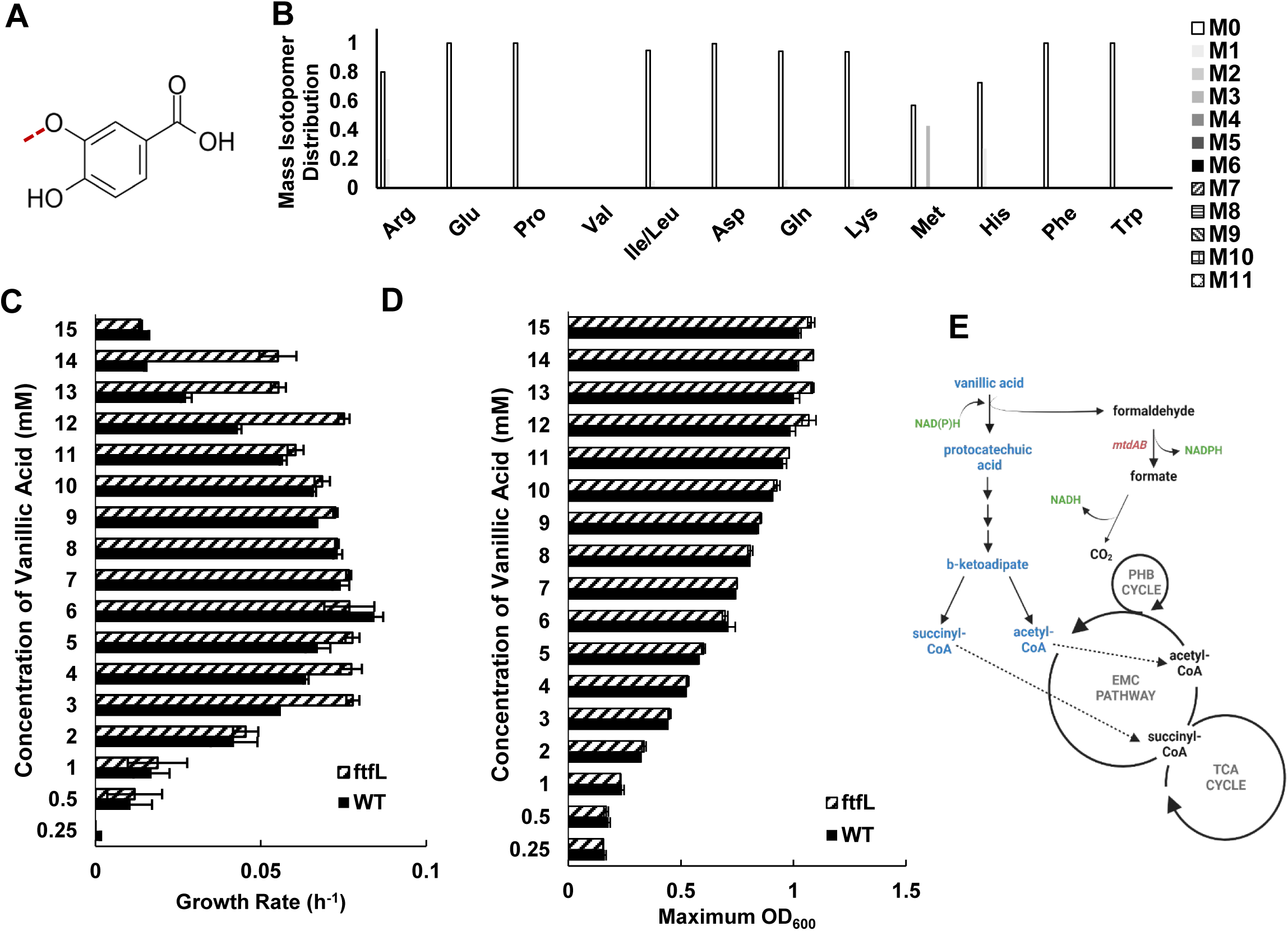
*M. extorquens* SLI 505 does not assimilate the formaldehyde released during the demethoxylation of vanillic acid into biomass. **A.** Vanillic acid with methoxy carbon labeled with ^13^C (indicated by red dashed line), hereafter referred to as ^13^C-vanillic acid. **B.** ^13^C fingerprinting of *M. extorquens* SLI 505 grown on 5 mM ^13^C-vanillic acid. M# indicates the number of ^13^C-carbons incorporated into the measured amino acid. **C**. Growth rates of wild-type *M. extorquens* SLI 505 (solid bar) and *M. extorquens* SLI 505 Δ*ftfL* (diagonal dashed bar) on 0.25-15 mM. Error bars represent standard deviation of two replicates. **D.** Maximum OD_600_ of wild-type *M. extorquens* SLI 505 (solid bar) and *M. extorquens* SLI 505 Δ*ftfL* (diagonal dashed bar) on 0.25-15 mM. Error bars represent standard deviation of two replicates. **E.** Updated metabolic map for growth on vanillic acid, eliminating formaldehyde assimilation as the metabolic fate of formaldehyde

The lack of contribution of formaldehyde towards biomass was also confirmed genetically. An essential gene for the assimilation of C_1_ intermediates into the serine cycle, *ftfL*^21^ (**Figure 1A)**, was deleted from *M. extorquens* SLI 505, and the mutant’s growth on aromatic acids was measured. The Δ*ftfL* strain has similar growth rates to wild-type on low and medium concentrations of vanillic acid and faster growth rates on high concentrations of vanillic acid (**Figure 3C**). Maximum OD_600_ values are consistent with wild-type values (**Figure 3D**). Based on labeling and growth phenotypes of mutants, we conclusively report that methoxydotrophic metabolism does not rely on the assimilation of formaldehyde (**Figure 3E**) and that biomass is therefore generated via heterotrophic assimilation of the aromatic acid ring carbons.

### Formaldehyde detoxification is beneficial but not necessary for the growth of methylotrophs on vanillic acid

The data thus far indicate that the fate of formaldehyde during growth of *M. extorquens* SLI 505 on vanillic acid is dissimilation to CO_2_. Here, an important distinction must be made between formaldehyde dissimilation versus detoxification. In methylotrophic bacteria, formaldehyde detoxification is necessarily coupled to formaldehyde dissimilation via the H_4_MPT-dependent formaldehyde oxidation pathway^21,23^. Formaldehyde oxidation to formate is coupled to the reduction of NADP^+^ to NADPH, and formate oxidation to CO_2_ can proceed via NAD-dependent or -independent formate dehydrogenases^21,23,24^. In summary, there are no metabolic routes for intracellular detoxification of formaldehyde that are not also linked to essential NADPH generation steps.

To disentangle these two processes, *mptG*, a gene involved in the biosynthesis of H_4_MPT involved in formaldehyde oxidation to fomate, was deleted in *M. extorquens* SLI 505; this mutant is unable to grow on C_1_ compounds^31^. Growth rates (**Figure 4A**) and maximum OD_600_ values (**Figure 4B**) for *M. extorquens* SLI 505 Δ*mptG* are reported. The Δ*mptG* strain has severely reduced growth rates at concentrations greater than 1 mM as compared to the wild-type strain yet still reaches the same maximum OD_600_ values. Measurements of intracellular formaldehyde concentrations of the Δ*mptG* strain (**Figure 4C**) reveal nearly 5-10x the amount of formaldehyde accumulation as compared to wild-type strain during growth on low and high concentrations of vanillic acid, consistent with the predicted phenotype of this mutant; the levels of formaldehyde accumulation in the Δ*mptG* strain are still well below what is considered toxic for *M. extorquens*.^30^ Thus, methylotrophic pathways for formaldehyde oxidation are important but not essential for growth on aromatic acids.

**Figure 4.**
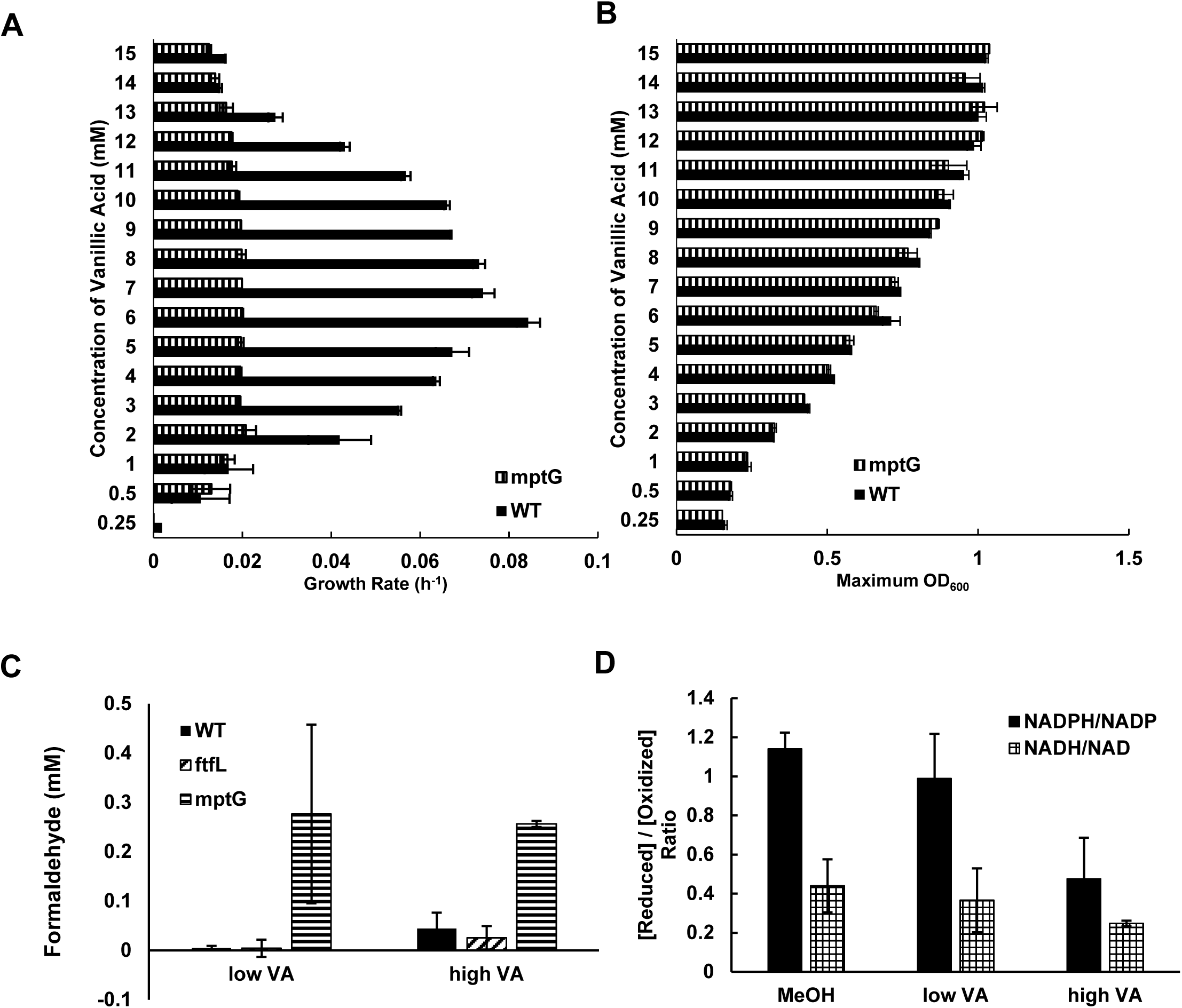
Identifying the relative contribution of formaldehyde detoxification towards growth on vanillic acid. **A.** Growth rates of wild-type *M. extorquens* SLI 505 (solid bar) and *M. extorquens* SLI 505 Δ*mptG* (vertical dashed bar) on 0.25-15 mM. Error bars represent standard deviation of two replicates. **B.** Maximum OD_600_ of wild-type *M. extorquens* SLI 505 (solid bar) and *M. extorquens* SLI 505 Δ*mptG* (vertical dashed bar) on 0.25-15 mM. Error bars represent standard deviation of two replicates. **C.** Intracellular formaldehyde concentrations of wild-type *M. extorquens* SLI 505 (solid bar), *M. extorquens* SLI 505 Δ*ftfL* (diagonal dashed bar), and *M. extorquens* SLI 505 Δ*mptG* (horizontal dashed bar) grown on “low” (5 mM) and “high” (12-13 mM) concentrations of vanillic acid. Error bars represent standard deviation of three biological replicates.**D.** Ratio of NADPH to NADP^+^ (solid bar) and NADH to NAD^+^ (checkered bar) during growth of wild-type *M. extorquens* SLI 505 on “low” (5 mM) and “high” (13 mM) concentrations of vanillic acid and 50 mM methanol. Error bars represent standard deviation of three biological replicates.

We questioned whether the growth rate defect of the Δ*mptG* strain was due to a buildup of formaldehyde (*i.e.* inefficient detoxification) or tied to energy conservation (*i.e.* formaldehyde dissimilation and NADPH production), both of which have been demonstrated to inhibit growth^24,32^. Canonical methylotrophic metabolism on substrates such as methanol is considered limited by NADPH, as NADPH must be spent for formate assimilation into the serine cycle and is generated from formaldehyde oxidation to formate ^24^. Several lines of evidence suggest that NADP(H) may also limit growth on vanillic acid by methylotrophs: (i) the initial demethoxylation of vanillic acid is coupled to NAD(P)H oxidation^29,33^, (ii) formaldehyde oxidation to formate follows methylotrophic pathways described above, (iii) growth on high concentrations of vanillic acid is correlated with high relative expression of genes involved in the linked EMC and polyhydroxybutyrate pathways at steps involving NADPH cycling, and growth on vanillic acid results in polyhydroxybutyrate accumulation, and (iv) genes for the NADPH-producing transhydrogenase *pntAB* are also highly expressed during growth on high concentrations of vanillic acid (**Supplementary Figure 1)**.

To identify reducing power limitations, if any, the ratio of NADPH to NADP^+^ and NADH to NAD^+^ during growth on low and high concentrations of vanillic acid was measured. Consistent with the predictions and preliminary data from above, the NADH/NAD^+^ ratios were consistent across vanillic acid concentrations but the NADPH/NADP^+^ ratios were lower at higher concentrations of vanillic acid and even lower than what is reported for methanol (**Figure 4D**). Thus, we concluded that growth on high concentrations of vanillic acid is at least partially limited by NADPH.

### Transcriptional response to growth on vanillic acid is concentration-dependent, mimicking heterotrophic growth at low concentrations and methylotrophic and heterotrophic growth at high concentrations

The strategy of formaldehyde dissimilation and heterotrophic assimilation during growth of *M. extorquens* SLI 505 on vanillic acid led us to investigate how operation of primary metabolic pathways differ during growth on vanillic acid compared to other substrates. RNA-seq was used to compare expression profiles of metabolic genes during growth on low (5 mM) and high (10 mM) vanillic acid, methanol, and acetate. Here, the methanol transcriptome is a baseline for the expression of metabolic genes during methylotrophic growth, and the acetate transcriptome is a baseline for the expression of metabolic genes during heterotrophic growth. **Figure 5** reports heatmaps for gene expression during aromatic acid metabolism and major methylotrophic and heterotrophic modules (see metabolic map, **Figure 1A**). To compare the relative expression of genes across four different substrate conditions, normalized Z-scores of genes are reported to identify common trends. Genes involved in aromatic acid catabolism are most highly expressed during growth on high concentrations of vanillic acid, to a lesser extent on low concentrations of vanillic acid, and not at all during growth on methanol or acetate, as is to be expected (**Figure 5A**). For both the low and high concentration of vanillic acid conditions, aromatic acid catabolism genes are some of the most highly expressed genes. Genes involved in methylotrophic pathways for formaldehyde oxidation, formate assimilation, and the serine cycle are most highly expressed during growth on methanol and least upregulated during growth on acetate (**Figure 5B, C, D, F**). The expression of these genes during growth on low concentrations of vanillic acid is at levels similar to what is observed on acetate. In contrast, expression levels on high concentrations of vanillic acid are closer to levels observed on methanol; this is surprising, considering methylotrophic assimilatory pathways are not utilized during growth on vanillic acid (**Figure 3**). Genes involved in heterotrophic pathways, such as the TCA cycle and the ethylmalonyl-CoA pathway used for glyoxylate regeneration, are also highly expressed during growth on high concentrations of vanillic acid. Overall, the data indicate that the expression patterns of metabolic genes on low concentrations of vanillic acid mimic those on acetate whereas expression patterns of metabolic genes on high concentrations of vanillic acid mimic those on methanol and acetate.

**Figure 5.**
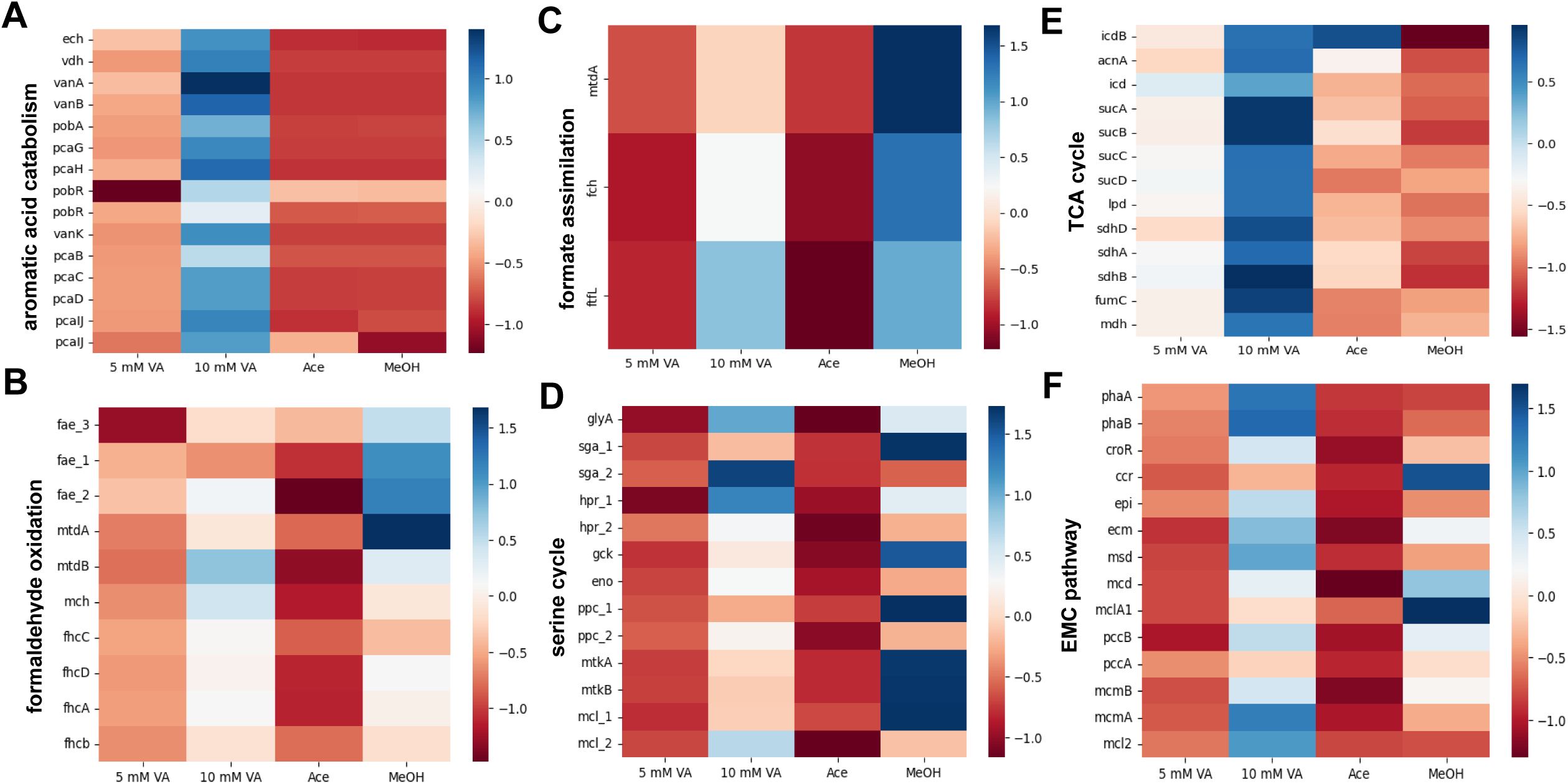
Normalized expression levels of methylotrophic and heterotrophic genes during growth on “low” (5 mM) and “high” (10 mM) concentrations of vanillic acid, 10 mM acetate, and 50 mM methanol. Expression profiles on methanol serve as a comparison for methylotrophic growth; expression profiles on acetate serve as a comparison for heterotrophic growth. Colors and corresponding color scale bar on the right of each graph indicates Z-score-corrected expression for comparisons of each gene across the four substrate conditions. Panels indicate metabolic modules relevant to methylotrophy and heterotrophy: **A**. aromatic acid gene island; **B**. H_4_MPT-dependent formaldehyde oxidation pathway; **C**. H_4_F-dependent formate assimilation pathway; **D**. serine cycle; **E**. TCA cycle; **F**. EMC pathway for glyoxylate regeneration

### Co-consumption of methanol and vanillic acid reveals two different strategies for the optimization of substrate consumption

Transcriptomic data from **Figure 5** indicates that vanillic acid induces expression of methylotrophic and heterotrophic metabolic pathways in *M. extorquens* SLI 505, yet we know that both cannot be^25^ and are not simultaneously employed in this organism (**Figure 3**). We chose to utilize a mixed substrate strategy to further interrogate the consequences of simultaneous expression of methylotrophic and heterotrophic pathways, by characterizing growth of *M. extorquens* SLI 505 on vanillic acid in combination with methanol. Because growth on vanillic acid has strong concentration-dependent phenotypes and different transcriptomic profiles, we investigated co-consumption patterns of *M. extorquens* SLI 505 at various vanillic acid concentrations. *M. extorquens* SLI 505 was grown on methanol with either a low concentration of vanillic acid (5 mM) or a high concentration of vanillic acid (13 mM). When *M. extorquens* SLI 505 was grown on low concentrations of vanillic acid with 50 mM methanol (**Figure 6A, B),** growth resulted in a diauxic shift with an initial growth phase that mimicked growth on methanol. We used ^13^C-vanillic acid and ^13^C-methanol to identify which substrates were allocated towards assimilation based on ^13^C fingerprinting analysis at various time points during growth. *M. extorquens* SLI 505 was grown on low concentrations of vanillic acid (5 mM), 50 mM methanol, and either 5 mM vanillic acid and 50 mM ^13^C-methanol (purple curve, **Figure 6A**) or 5 mM ^13^C-vanillic acid and 50 mM methanol (purple curve, **Figure 6B**). Cells were harvested at the time points indicated by arrows that correspond with mid- to late-exponential growth of each diauxic growth phase, and amino acids were analyzed for the presence or absence of ^13^C to correlate with the assimilation of labeled substrate. **Figures 6C** and **6D** show the mass isotopomer distribution for proteinogenic amino acids harvested at different time points during growth on unlabeled vanillic acid and labeled methanol. ^13^C fingerprinting reveals the presence of heavily labeled amino acids during time point 1, consistent with our hypothesis that methanol is used first. At time point 2 (**Figure 6D**), some amino acids are labeled but there is a reduction from timepoint 1, likely due to a switch to an unlabeled substrate (vanillic acid) with carryover of previously-labeled intermediates from growth on methanol. ^13^C fingerprinting results from the converse experiment (**Figure 6B**) also agree with our hypothesis that methanol is preferentially utilized over low concentrations of vanillic acid. **Figure 6E** and **6F** show the mass isotopomer distribution for proteinogenic amino acids harvested at different time points during growth on labeled vanillic acid and unlabeled methanol; both time points show no label incorporation into amino acids, indicating that the methoxy-carbon of vanillic acid is not assimilated during either stage of growth. Nucleotide ratios mimic patterns shown on either substrate alone, with an increase in NADH/NAD^+^ that could be due to increased flux from formate to CO_2_ via NAD-dependent formate dehydrogenases (**Figure 6G**).

**Figure 6.**
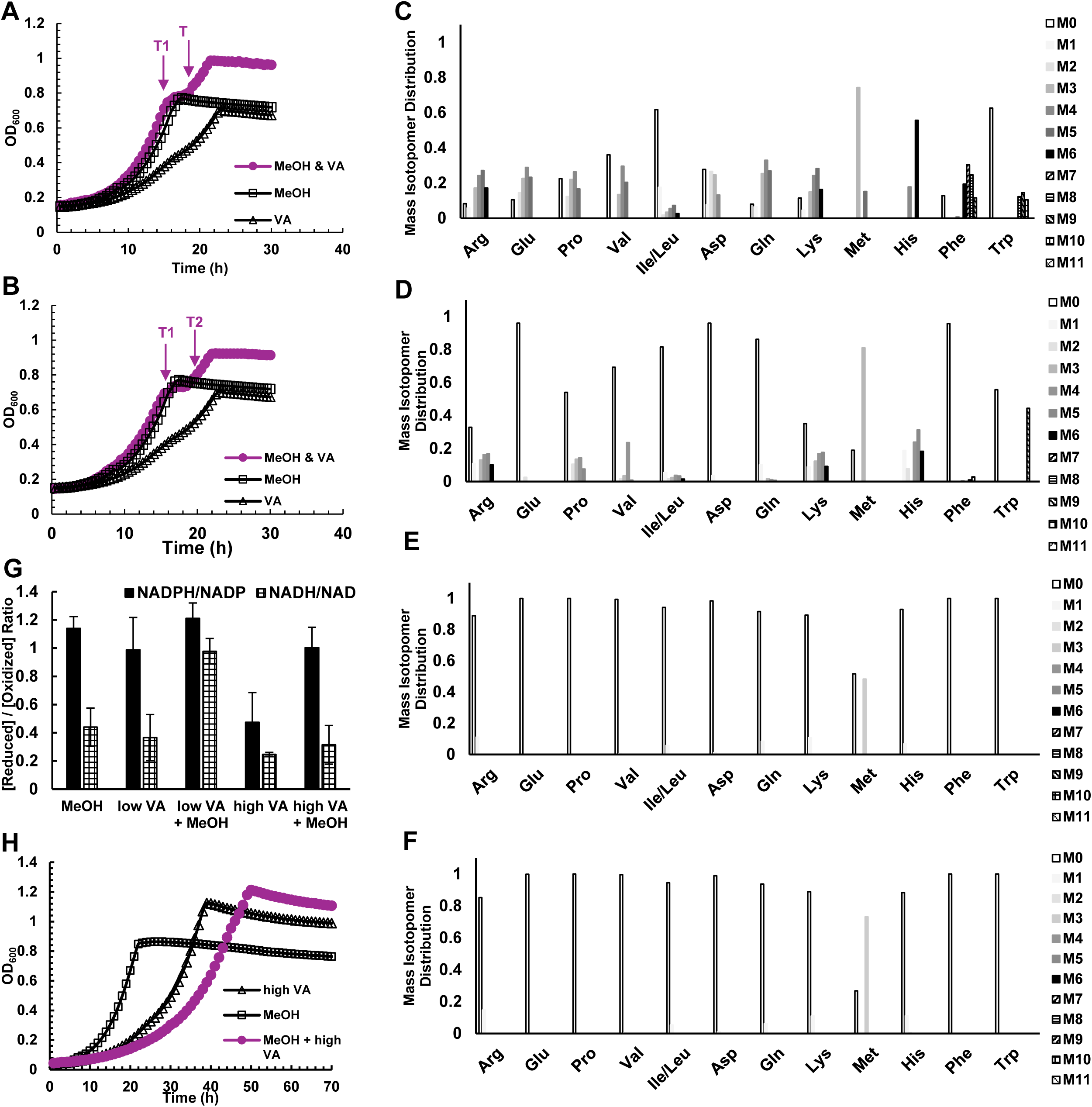
Dual-substrate growth of *M. extorquens* SLI 505 on vanillic acid and methanol. **A.** Growth curves of *M. extorquens* SLI 505 on 5 mM vanillic acid (“VA”, open triangles), 50 mM methanol (“MeOH”, open squares), and 5 mM vanillic acid + 50 mM ^13^C-methanol (“MeOH & VA”, closed purples circles). Arrows indicates time points (T1 and T2) at which samples were harvested for ^13^C-fingerprinting analysis. **B.** Growth curves of *M. extorquens* SLI 505 on 5 mM vanillic acid (“VA”, open triangles), 50 mM methanol (“MeOH”, open squares), and 5 mM ^13^C-vanillic acid + 50 mM methanol (“MeOH & VA”, closed purple circles). Arrows indicates time points (T1 and T2) at which samples were harvested for ^13^C-fingerprinting analysis. **C.** ^13^C fingerprinting of time point 1 (T1) of *M. extorquens* SLI 505 grown on 5 mM vanillic acid + 50 mM ^13^C-methanol. M# indicates the number of ^13^C-carbons incorporated into the measured amino acid. **D.** ^13^C fingerprinting of time point 1 (T2) of *M. extorquens* SLI 505 grown on 5 mM vanillic acid + 50 mM ^13^C-methanol. M# indicates the number of ^13^C-carbons incorporated into the measured amino acid. **E.** ^13^C fingerprinting of time point 1 (T1) of *M. extorquens* SLI 505 grown on 5 mM ^13^C-vanillic acid + 50 mM methanol. M# indicates the number of ^13^C-carbons incorporated into the measured amino acid. **F.** ^13^C fingerprinting of time point 1 (T2) of *M. extorquens* SLI 505 grown on 5 mM ^13^C-vanillic acid + 50 mM methanol. M# indicates the number of ^13^C-carbons incorporated into the measured amino acid. **G.** Ratio of NADPH to NADP^+^ (solid bar) and NADH to NAD^+^ (checkered bar) during growth of *M. extorquens* SLI 505 on 50 mM methanol (MeOH), 5 mM vanillic acid (low VA), 5 mM vanillic acid + 50 mM methanol (low VA + MeOH). Error bars represent standard deviation of three biological replicates. **H.** Growth curves of *M. extorquens* SLI 505 on 13 mM vanillic acid (“high VA”, open triangles), 50 mM methanol (“MeOH”, open squares), and 13 mM vanillic acid + 50 mM methanol (“MeOH & high VA”, closed purple circles)

To investigate if these consumption patterns are similar at high concentrations of vanillic acid, *M. extorquens* SLI 505 was grown on 13 mM vanillic acid with 50 mM methanol and compared to growth on either high vanillic acid or methanol alone **(Figure 6H**). Interestingly, growth on dual substrates had longer lag times than on either substrate alone, in sharp contrast to what was observed during growth on low concentrations of vanillic acid and methanol. Growth on high concentrations of vanillic acid with methanol restores the NADPH/NADP^+^ and NADH/NAD^+^ ratios to what is observed during growth on methanol or low concentrations of vanillic acid (**Figure 6G**) and diauxic growth is no longer observed (**Figure 6H**).

## Discussion

How complex substrates are converted to various intermediates that can be assimilated through two drastically different modes of metabolism is a fundamental gap in our understanding of microbial physiology^34,35^. Aromatic acid metabolism is a model by which to understand how methylotrophic bacteria maximize bioenergetic yields during growth on multiple intermediates and can bolster our understanding of how facultative methylotrophs balance methylotrophy and heterotrophy. Here, we establish *M. extorquens* SLI 505 as a model organism for vanillic acid metabolism. Genes involved in methoxylated aromatic acid catabolism are reported and shown to be organized similarly to what has been found in other *extorquens* clade members capable of aromatic acid metabolism. Growth phenotypes on a wide range of vanillic acid concentrations are starkly concentration-dependent, with reduced growth at higher substrate concentrations that we hypothesize is related to NADPH limitation rather than formaldehyde accumulation and mixed utilization patterns when grown on vanillic acid with methanol.

To date, much of the literature surrounding aerobic aromatic acid metabolism in bacteria has focused on mechanisms by which aromatic rings are converted into building blocks for assimilation and how bacteria can salvage carbon from complex substrates^8,9,9,36^. The toxicity of aromatic acids, whether due to their inherent chemical properties or C_1_ intermediates that arise in their degradation, has been considered a biochemical inevitability during the metabolism of these compounds and/or a target for metabolic engineering for biotechnological applications^12,18^. However, the recent focus on the ability of particular clades of methylotrophic bacteria to robustly grow on aromatic acids has expanded our understanding of how lignin-derived aromatic compounds might influence microbial physiology in natural environments and how this might be co-opted for biotechnology^19,26,37^. Methylotrophic bacteria are especially attractive organisms for investigating aromatic acid metabolism because they have evolved elegant mechanisms by which not only to detoxify but also assimilate high concentrations of formaldehyde as a routine consequence of their metabolism^21,23^. Non-methylotrophic bacteria that natively encode formaldehyde detoxification mechanisms, such as *Pseudomonas putida* KT2440, display a marked decrease in growth due to formaldehyde accumulation even at low concentrations of vanillic acid and efforts to engineer *P. putida* KT2440 to overcome this limitation are ongoing^18,38^. In contrast, *M. extorquens* SLI 505 is an environmental isolate naturally capable of robust aromatic acid metabolism without concomitant formaldehyde accumulation, even during growth on concentrations as high as 12 mM (**Table 1**).

^13^C fingerprinting with methoxy-labeled ^13^C-vanillic acid (**Figure 3A**) was used to track labeled carbon in amino acids as a proxy for determining whether formaldehyde generated from vanillic acid was assimilated towards biomass. Of the detectable proteinogenic amino acids, labeled carbon was absent from all except for methionine, which was roughly 40% labeled (**Figure 3B)**. Methionine biosynthesis requires abstraction of methyl groups and the partial label incorporation is likely a reflection of methionine biosynthesis alone and not indicative of assimilation of formaldehyde, as labeled carbon is absent from all other amino acids. Other studies using labeled vanillic acid for investigating aromatic acid metabolism in the non-methylotrophic bacterium, *Sphingobium sp.* SYK-6 also noted labeling of methionine and links between methionine and C_1_ metabolism^39^. Despite the minor labeling of methionine, we still conclude that formaldehyde generated during growth on vanillic acid is not assimilated towards biomass.

When methylotrophic bacteria grow on methanol, 100% of the initial substrate must be converted to formaldehyde before it is shunted towards assimilatory or dissimilatory pathways^21,24^. In contrast, the growth of methylotrophic bacteria on methoxylated aromatic acids mainly produces heterotrophic intermediates and only results in an eighth of the total carbon in the substrate being converted to formaldehyde (**Figure 1A, 3E**). Thus, it is not surprising that formaldehyde toxicity is not a major constraint during this metabolism based solely on differences in the relative amounts of formaldehyde generated by vanillic acid. Although we hypothesize that the more substantial constraints on metabolism are at the level of currency metabolites, we cannot rule out the possibility that other metabolic or non-metabolic processes contribute to the concentration-dependent phenotypes we report here. We see no evidence for stress response in our transcriptomic data set for high vanillic acid concentrations in comparison to other substrate conditions, and the growth media is sufficiently buffered to prevent vanillic acid itself from causing acute acid stress. A recent study suggested that *M. extorquens* experiences membrane depolarization as a result of excess vanillic acid diffusion across membranes that can lead to disruption of proton motive force for energy production^37^. Our preliminary analysis of membrane permeability (**Supplementary Figure 2**) of *M. extorquens* SLI 505 in a variety of substrates and vanillic acid concentrations did not indicate a correlation with growth on high concentrations of vanillic acid. However, a link between membrane depolarization and energy metabolism could relate to the lack of growth of *M. extorquens* SLI 505 at vanillic acid concentrations higher than 15 mM and warrants further investigation in this system.

The fate of formaldehyde in methylotrophic bacteria is unique in that formaldehyde detoxification is necessarily coupled to formaldehyde dissimilation. In literature discussing methylotrophy, the terms detoxification and dissimilation are often used interchangeably as they relate to this metabolism. Here, we take advantage of mutants deficient in formaldehyde assimilation (Δ*ftfL*) and formaldehyde detoxification and dissimilation (Δ*mptG*) to disentangle these processes. Because aromatic acid metabolism does not require formaldehyde assimilation for growth, it is a useful model to interrogate these processes. *M. extorquens* SLI 505 Δ*ftfL* is incapable of assimilating formaldehyde yet does not accumulate formaldehyde (**Figure 4C**) and grows faster than wild-type at most vanillic acid concentrations (**Figure 3C**). The latter was a surprising growth phenotype, as loss of *ftfL* has no obvious advantage if assimilation of C_1_ units does not naturally occur during this metabolism. Growth advantages due to loss of *ftfL* have been reported in *M. extorquens* AM1 evolved in succinate^40^, but physiological benefits during growth on vanillic acid remain unknown.

In contrast, a Δ*mptG* mutant incapable of synthesizing the carbon carrier necessary for formaldehyde oxidation to formate has substantially slower growth rates than wild-type at all vanillic acid concentrations (**Figure 4A**) yet manages to reach the same final OD_600_ values as wild-type (**Figure 4B**). Interestingly, addition of lanthanum chloride significantly improves growth rates of *M. extorquens* SLI 505 Δ*mptG* during growth on vanillic acid (data not shown). We hypothesize that this is due to lanthanum chloride inducing expression of lanthanide-dependent alcohol dehydrogenases that might play a role in formaldehyde oxidation,^41^ and investigations into the role of lanthanides during growth on vanillic acid is the subject of ongoing work. Growth phenotypes of *M. extorquens* SLI 505 Δ*mptG* sharply contrast what has recently been reported for a strain of *M. extorquens* PA1 engineered to metabolize vanillic acid, where a Δ*mptG* mutation is fatal^37^. The differences in the essentiality of *mptG* in our natural isolate *M. extorquens* SLI 505 versus the engineered *M. extorquens* PA1 strain could highlight differences in evolution or metabolic regulation of formaldehyde-related processes in these two systems that are worth investigating further. Here, we hypothesized that the growth rate defect of our Δ*mptG* strain was not due to formaldehyde accumulation, as intracellular formaldehyde levels of this mutant were below toxic levels for this strain (**Figure 4C)**^30^, but rather due to the importance of formaldehyde dissimilation and nucleotide pools in the form of NADP^+^ and NADPH. This is not unprecedented, as tight regulation between NAD(P)H pools, cofactor selectivity, and vanillic acid metabolism have been reported in *Pseudomonas putida* KT2440^29^ and *Sphingobium sp.* SYK-6^39^. Methylotrophic metabolism has been shown to be NADPH-limited^24^, and this appears to hold true even for substrates such as vanillic acid that do not require formaldehyde assimilation. Future work will investigate nucleotide pools in Δ*mptG* strains to further substantiate our hypothesis about NADPH limitation generating a partial bottleneck for carbon assimilation during growth on high concentrations of vanillic acid.

Growth on methanol and low concentrations of vanillic acid alleviates some of the growth phenotypes exhibited during growth on low concentration of vanillic acid alone (**Figure 6A, B**). Several important conclusions can be drawn about how *M. extorquens* SLI 505 copes with growth on low concentrations of vanillic acid with methanol: (i) consumption follows a hierarchy, where growth on methanol occurs before growth on vanillic acid with a marked diauxic shift, (ii) growth on vanillic acid in the second phase of the growth curve occurs without the lag characteristic of growth on vanillic acid alone, (iii) methanol itself induces expression of methylotrophic pathways^42^, yet despite both methylotrophic and heterotrophic pathways being operational, vanillic acid is still metabolized as shown in **Figure 4E**. Here, the presence of methanol with low concentrations of vanillic acid would theoretically “prime” the cells for methylotrophy; this would shift the expression profile in this condition to reflect growth on high concentrations of vanillic acid, where both methylotrophic and heterotrophic pathways are expressed (**Figure 5**). Yet, growth phenotypes do not necessarily correlate with predictions made from transcriptomic analyses alone, as diauxic growth as a result of operation of both methylotrophic and heterotrophic pathways does not occur during growth on methanol and high concentrations of vanillic acid (**Figure 6H**). The vanillic acid used in this experiment is not fully labeled. Thus we cannot rule out the possibility that the unlabeled ring carbons of vanillic acid is being metabolized and/or assimilated in combination with methanol during the first stage of growth. However, this scenario would require simultaneously operating methylotrophic and heterotrophic pathways which is unlikely to occur.

Traditionally, sequential utilization of substrates occurs if a preferred substrate supports a higher growth rate, assuming both substrates are in excess^34^. Limitation of substrates eliminates sequential or preferential substrate utilization in favor of co-utilization to maximize growth^35,43^. In contrast to this paradigm, growth on methanol and high concentrations of vanillic acid exacerbates the growth phenotypes exhibited during growth on high concentrations of vanillic acid alone (**Figure 6H**). Further labeling and ^13^C fingerprinting studies will be required to validate if a single or both substrates are being consumed but catabolic repression via diauxic growth is no longer present.

In recent decades, methylotrophs have emerged as promising model organisms for biotechnological manipulation due to their genetic tractability, multi-omics characterizations, and demonstrated flux through pathways directly linked to the production of value-added chemicals^20,44^. However, much of this work has been done with renewable C_1_ feedstocks such as methane or methanol^20,45–47^. There are vast biotechnological implications for robust aromatic acid metabolism in methylotrophs that does not result in formaldehyde accumulation^3,7,48–50^. We report growth of *M. extorquens* SLI 505 on vanillic acid concentrations as high as 15 mM, substantially higher than what has been reported by many other model organisms with well-characterized aromatic acid metabolisms^9,39^. It has been shown that the inclusion of *M. extorquens* PA1 (non-aromatic acid utilizer) in a lignin-degrading consortium is sufficient to detoxify formaldehyde^17^; similar studies have not been replicated for *M. extorquens* strains naturally capable of aromatic acid metabolism. Additionally, we have demonstrated that growth on high concentrations of vanillic acid is natively coupled to the accumulation of the bioplastic polyhydroxybutyrate (**Supplementary Figure 1C**), providing an attractive starting point for future metabolic engineering efforts.

## Materials and Methods

### Bacterial strains and cultivation

Isolation of *Methylobacterium extorquens* SLI 505 is described previously^26^. *M. extorquens* SLI 505 (wild-type and mutant strains) was grown in minimally defined *Methylobacterium* PIPES (MP) media^51^ supplemented with exogenous carbon sources as indicated. Vanillic acid was prepared fresh every time by dissolving vanillic acid powder (Sigma Aldrich) to the desired final concentration (0.25-20 mM) in sterile MP with a sterile stir-bar. Vanillic acid was not soluble in MP beyond 20 mM. 3 mL pre-cultures of *M. extorquens* SLI 505 in MP with 15 mM succinate (Millipore Sigma) were grown overnight in 14-mL polystyrene round-bottom plastic culture tubes (Falcon) at 30°C and 200 rpm in an Innova S44i incubator shaker (Eppendorf). Pre-cultures were washed twice in MP by centrifugation at 2000 x g for 10 minutes and diluted into a desired volume of fresh vanillic acid media in MP to an OD600 of 0.1. Growth curves in **Figure 2**, **Figure 3C**, **Figure 3D**, **Figure 4A**, and **Figure 4B** were generated by growing *M. extorquens* SLI 505 (wild-type or mutants) at the indicated substrate concentrations as 650 µL cultures in transparent 48-well plates (Corning) incubated at 30°C with orbital shaking at 548 rpm with OD_600_ readings every 30 minutes using a Synergy HTX plate reader (Biotek). Experimental conditions were run in duplicates and each growth curve experiment was independently run twice to validate growth phenotypes, for a total of four replicates per condition. Only two replicates from a single run are reported, and >10 exponential-phase data points were used for linear regression analysis to calculate growth rates.

### Cluster analysis for aromatic acid gene island visualization

The genomes of *M. extorquens* SLI 505 and *Methylobacterium sp.* AMS5 were downloaded from Integrated Microbial Genomes Joint Genome Institute and NCBI in January 2025. The aromatic acid island region in *M. sp.* AMS5 was identified previously^19^, spanning from locus tag Y590_RS18530 to Y590_RS18605. Homologous regions were identified in *M. extorquens* SLI 505 spanning from locus tag JSW75_000124 to JSW75_000140. Genomic regions were aligned via Clinker CAGECAT using default settings^27^.

### DNA manipulation

All strains and plasmids used in this study are listed in Supplementary Table 1. Deletion plasmids were made using a suicide vector, pCM433KanT, with *sacB* counterselection marker^52^. Fragments for deletion vectors were constructed by inserting 500-800 bp regions of *M. extorquens* SLI 505 genomic DNA flanking the gene of interest into pCM433KanT via HiFi DNA Assembly MasterMix (New England Biolabs). Following full-plasmid sequence verification (Plasmidsaurus), deletion plasmids were transformed into *Escherichia coli* S17. Biparental mating was performed between *E. coli* S17 containing the plasmid of interest *M. extorquens* SLI 505. *M. extorquens* SLI 505 with an integrated plasmid (single crossover) was selected for growth on minimal agar plates supplemented with 15 mM succinate, 50 ug/mL kanamycin, and methylamine as the nitrogen source to prevent growth of donor *E. coli* S17 strains. Counterselection (double crossover) was performed on minimal media agar plates with 15 mM succinate and 10% sucrose to identify colonies that have lost the plasmid. Replica patching colonies onto minimal agar plates containing succinate and kanamycin and succinate and sucrose allowed for selection of candidates capable of growth on sucrose and incapable or growth on kanamycin, and candidates were PCR-screened and sequenced to confirm deletions of the gene of interest.

### RNA sequencing

Pre-cultures of *M. extorquens* SLI 505 were grown and prepared as described above and used to inoculate triplicate 250-mL glass culture flasks containing 50 mL of MP supplemented with either 5 mM vanillic acid, 10 mM vanillic acid, 50 mM MeOH, or 10 mM acetate at an OD_600_ of 0.1. 40 mL of culture were harvested at mid-exponential phase (OD_600_ of 0.5-1 for all conditions) by centrifugation at 4°C at 2000 x g for 10 minutes. Total RNA was extracted using the Qiagen RNeasy Kit and quantified. rRNA depletion using the Ribo-Zero RNA Plus rRNA Depletion Kit (Illumina), library preparation, and Illumina Hi-Seq sequencing was performed by SeqCoast Genomics. KBase was used to align 12 million 2 x 150 bp paired-end reads per sample via HISAT2 to *M. extorquens* SLI 505 genome from NCBI, assemble transcripts via StringTie, and call differentially expressed genes via DESeq2^53^. Z-scores normalized to total counts are reported in **Figure 5** and **Supplementary Figure 1** to facilitate comparisons across four substrate conditions.

### Measurement of intracellular formaldehyde concentrations

Pre-cultures of *M. extorquens* SLI 505, Δ*ftfL*, and/or Δ*mptG* were grown and prepared as described above and used to inoculate triplicate 250-mL glass culture flasks containing 50 mL of MP supplemented with 5-15 mM vanillic acid, as indicated, at an OD_600_ of 0.1. 40 mL of culture was harvested at mid-exponential phase (OD_600_ of 0.5-1 for all conditions) and lysed at 25 kpsi using a MC-BA Cell Disruptor (Constant Systems). Lysates were kept on ice and processed immediately. Intracellular formaldehyde concentrations from cell lysates were determined using a colorimetric Purpald (4-amino-3-hydrazino-5-mercapto-1,2,4-triazole)-based assay^54^, as previously described^41^.

### Measurement of nucleotide pools

Pre-cultures of *M. extorquens* SLI 505 were grown and prepared as described above and used to inoculate triplicate 14-mL polystyrene round-bottom plastic culture tubes (Falcon) containing 3 mL of MP supplemented with 5-13 mM vanillic acid, as indicated, at an OD_600_ of 0.1. 3 mL of culture was harvested at mid-exponential phase (OD_600_ of 0.5-1 for all conditions) and lysed at 25 kpsi using a MC-BA Cell Disruptor (Constant Systems). Lysates were kept on ice and processed immediately. Individual NAD+, NADP, NADH, and NADPH concentrations were measured using the Promega Bioluminescent Glo Assay, per published kit instructions.

### Measurement of relative polyhydroxybutyrate concentrations

Pre-cultures of *M. extorquens* SLI 505 were grown and prepared as described above and used to inoculate triplicate 14-mL polystyrene round-bottom plastic culture tubes (Falcon) containing 3 mL of MP supplemented with either 50 mM MeOH, 5 mM vanillic acid, or 10 mM vanillic acid at an OD_600_ of 0.1. 500 µL of culture were harvested at mid-exponental phase (OD_600_ of 0.5-1 for all conditions) and centrifuged at maximum speed for 5 minutes. Supernatants were decanted and cell pellets were resuspended in 20 µL of 0.5% Nile Blue A (Sigma Aldrich), then incubated at room temperature for 10 minutes. 200 µL of MP was added to each resuspension, and samples are centrifuged at maximum speed for 2 minutes. Supernatants were decanted and cell pellets were resuspended in 200 µL of 8% acetic acid, then incubated at room temperature for 1 minute. Samples were centrifuged at maximum speed for 1 minute, supernatant was decanted, and cell pellets were resuspended in 250 µL of MP. 200 µL of each sample were transferred to a black/clear bottom 96-well plate (Corning) and fluorescence was measured using a Spectramax plate reader with excitation at 510 nm and emission at 590 nm. Fluorescence values were normalized by subtracting the fluorescence of MP alone, then dividing relative fluorescence (RFU) by OD_600_ at time of harvesting and mM carbon to normalize across substrates.

### Measurement of membrane permeability using propidium iodide

Pre-cultures of *M. extorquens* SLI 505 were grown and prepared as described above and used to inoculate duplicate 14-mL polystyrene round-bottom plastic culture tubes (Falcon) containing 3 mL of MP supplemented with 15 mM succinate, 50 mM methanol, 10 mM acetate, 5 mM vanillic acid, 10 mM vanillic acid, or 13 mM vanillic acid. Cultures were harvested mid-exponential phase for all conditions (OD_600_ of 0.5-1 for all conditions), and two identical dilutions of each culture to an OD_600_ of 0.5 in 1 mL of MP were prepared. Samples were centrifuged at 2000 x g for 1 minute. Supernatants were discarded and cell pellets were resuspended in 1 mL of MP. Triplicate aliquots of 150 µL of each sample were transferred to a black/clear bottom 96-well plate (Corning) to serve as a background fluorescence correction of unstained cells. To the remaining 550 µL of each sample, propidium iodide (Thermo Fisher) was added to a final concentration of 5 µg/mL. Stained cells were incubated in the dark for 15 minutes. Triplicate aliquots of 150 µL of each sample were transferred to a black/clear bottom 96-well plate (Corning). Fluorescence was measured using a Spectramx plate reader with excitation at 493 nm and emission at 636 nm for detection of propidium iodide unbound to DNA (low membrane permeability) and with excitation at 535 nm and emission at 617 nm for detection of propidium iodide bound to DNA (high membrane permeability). Average corrected propidium iodide-bound measurements are reported.

### ^13^C fingerprinting and LC-MS analysis for tracking labeled carbon in amino acid

Vanillic acid with ^13^C methoxy carbon (referred to as ^13^C-vanillic acid) was synthesized by Sigma Aldrich. Pre-cultures of *M. extorquens* SLI 505 were grown and prepared as described above and used to inoculate triplicate 14-mL polystyrene round-bottom plastic culture tubes containing 4 mL of MP supplemented with 5 mM ^13^C-vanillic acid, 5 mM vanillic acid, 5 mM ^13^C-vanillic acid and 50 mM MeOH, or 5 mM vanillic acid and 50 mM ^13^C-MeOH (Sigma Aldrich), as indicated. 4 mL of culture were harvested at mid-exponential phase via fast-filtration using Nyaflo 0.2 µM 47 mM nylon filters (Pall Corporation) and filters containing cell biomass were immediately transferred to 50-mL tubes and flash frozen with liquid nitrogen. 8 mL of boiling sterile ultrapure water were added to each filter and boiled for 8 minutes. Samples were immediately cooled on ice for approximately 5 minutes, or until samples equilibrated to room temperature. The filters were removed from the tubes and the liquid was filter-sterilized using 0.22 µM filters into new tubes, and flash frozen with liquid nitrogen. Samples were completely lyophilized and reconstituted in 30:70 water:acetonitrile immediately prior to mass spectrometry analysis. Amino acid samples were analyzed using a liquid chromatography (LC) system (1200 series, Agilent Technologies, Santa Clara, CA) that was connected in line with an LTQ-Orbitrap-XL mass spectrometer equipped with an electrospray ionization (ESI) source (Thermo Fisher Scientific, Waltham, MA). The instrumentation is located in the QB3/Chemistry Mass Spectrometry Facility at the University of California, Berkeley. The LC system was equipped with a G1322A solvent degasser, G1311A quaternary pump, G1316A thermostatted column compartment, and G1329A autosampler unit (Agilent). The column compartment was equipped with an XBridge hydrophilic interaction liquid chromatography (HILIC) column (length: 100 mm, inner diameter: 2.1 mm, particle size: 3.5 micrometers, part number 186004433, Waters, Milford, MA). Ammonium formate (99%, Alfa Aesar, Ward Hill, MA), acetonitrile, formic acid (Optima LC-MS grade, 99.9% minimum, Fisher, Pittsburgh, PA), and water purified to a resistivity of 18.2 MΩ·cm (at 25 °C) using a Milli-Q Gradient ultrapure water purification system (Millipore, Billerica, MA) were used to prepare mobile phase solvents. Mobile phase solvent A was water and mobile phase solvent B was 90% acetonitrile/10% water, both of which contained 10 mM ammonium formate and 0.1% formic acid (volume/volume). The elution program consisted of isocratic flow at 99.5% (volume/volume) B for 2 min, a linear gradient to 40% B over 8 min, isocratic flow at 40% B for 3 min, a linear gradient to 99.5% B over 1 min, and isocratic flow at 99.5% B for 16 min, at a flow rate of 300 µL/min. The column compartment was maintained at 30 °C and the sample injection volume was 20 µL. External mass calibration was performed in the positive ion mode using the Pierce LTQ ESI positive ion calibration solution (catalog number 88322, Thermo Fisher Scientific). Full-scan, high-resolution mass spectra were acquired in the positive ion mode over the range of mass-to-charge ratio (*m*/*z*) = 65 to 250, using the Orbitrap mass analyzer, in profile format, with a mass resolution setting of 60,000 (at *m*/*z* = 400, measured at full width at half-maximum peak height, FWHM). Data acquisition and analysis were performed using Xcalibur software (version 2.0.7, Thermo).

## Acknowledgements

We thank all members of the Martinez-Gomez Lab at the University of California, Berkeley for their thoughtful suggestions during the experiment planning and writing stages of this manuscript. Thank you to Dr. Anthony T. Iavarone and the staff at the QB3/Chemistry Mass Spectrometry Facility at the University of California, Berkeley for valuable expertise and running of all mass spectrometry samples. The QB3/Chemistry Mass Spectrometry Facility received National Institutes of Health support (grant number 1S10OD020062-01). The information, data, or work presented herein was funded by the United States Department of Energy, Office of Science, Office of Biological and Environmental Research, under Award Number DE-SC0022318 subaward SH5849-772894; by the National Science Foundation under Grant 2142154; by the National Science Foundation under Grant 2127732. A.M.G. was supported in part by the National Institutes of Health Genetic Dissection of Cells and Organisms Training Grant 1T32GM132022-01.

