## Supplementary Information for "Aromatic acid metabolism in *Methylobacterium extorquens* reveals interplay between methylotrophic and heterotrophic pathways"

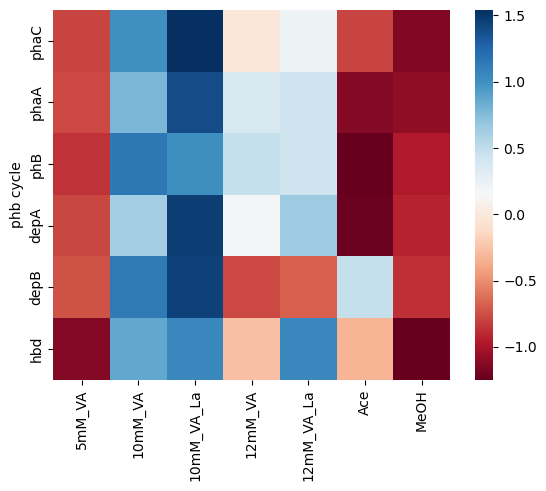

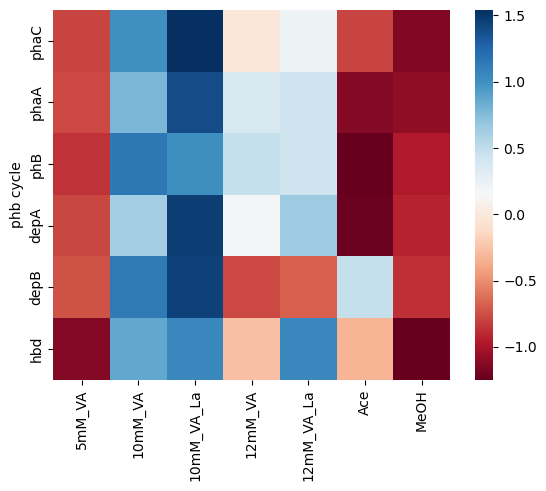

**PHB cycle genes**

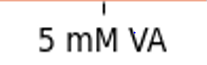

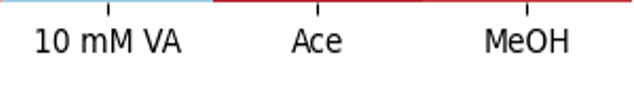

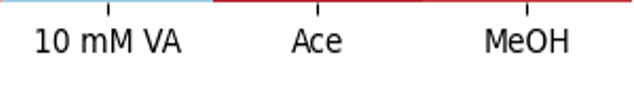

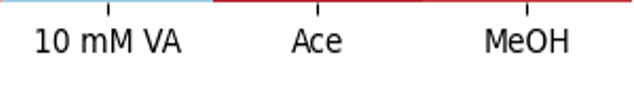

**C**

**A**

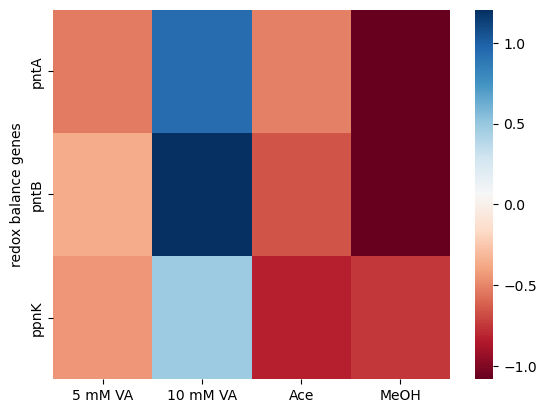

**redox balance genes**

**A**

**A**

**B**

**A**

|  | RFU/OD600/mM carbon |
| --- | --- |
| 5 mM VA | 10.3609 |
| 10 mM VA | 34.9112 |
| MeOH | 11.1051 |

**Supplementary Figure 1.** Evidence for NADPH as an important currency metabolite during growth on vanillic acid. Normalized expression levels of methylotrophic and heterotrophic genes during growth on “low” (5 mM) and “high” (10 mM) concentrations of vanillic acid, 10 mM acetate, and 50 mM methanol. Expression profiles on methanol serve as a comparison for methylotrophic growth; expression profiles on acetate serve as a comparison for heterotrophic growth. Colors and corresponding color scale bar on the right of each graph indicates Z-score-corrected expression for comparisons of each gene across the four substrate conditions. **A.** polydhydroxybutyrate (PHB) cycle genes; **B.** redox balance genes. **C.** Nile Blue A staining and normalized fluorescent measurements of *M. extorquens* SLI 505 in low vanillic acid (5 mM VA), high vanillic acid (10 mM VA), and 50 mM methanol as a relative measurement of polyhydroxybutyrate concentrations. Averages of three biological replicates are reported. Error bars represent standard deviation

|  | Normalized fluorescence of propidium iodide bound to DNA |
| --- | --- |
| succinate | 304180 |
| methanol | 364979 |
| acetate | 391884 |
| 5 mM VA | 396096 |
| 10 mM VA | 388019 |
| 13 mM VA | 340638 |

**Supplementary Figure 2.** Propidium iodide staining of *M. extorquens* SLI 505 as a measurement of membrane permeability. *M. extorquens* SLI 505 was grown in 15 mM succinate (“succinate”), 50 mM methanol (“methanol”), 10 mM acetate (“acetate”), 5 mM vanillic acid (“5 mM VA”), 10 mM vanillic acid (“10 mM VA”), and 13 mM vanillic acid (“13 mM VA”) and stained with propidium iodide, a dye that fluoresces at 535/617 nm upon binding of DNA and can act as a proxy for measuring membrane permeability. Averages of two biological replicates and three technical replicates of samples corrected for background fluorescence of unstained cells are reported. Error bars represent standard deviation

**Supplementary Table 1.** Strains and plasmids used in this study

| **Strain or Plasmid** | **Description** | **Reference** |
| --- | --- | --- |
| **Strains** |  |  |
| *Escherichia coli* 10β | Electrocompetent cloning strain | Invitrogen |
| *E. coli* S17-1 | Conjugating donor strain | New England Biolabs |
| *Methylobacterium extorquens* SLI 505 | Wild-type | 26 |
| *M. extorquens* SLI 505 Δ*ftfL* | Δ*ftfL* | This study |
| *M. extorquens* SLI 505 Δ*mptG* | Δ*mptG* | This study |
| **Plasmids** |  |  |
| pCM433KanT | SacB allelic exchange backbone; Km^R^ | 51 |
| pAG1 | SacB allelic exchange *ftfL* donor; Km^R^ | This study |
| pAG24 | SacB allelic exchange *mptG* donor; Km^R^ | This study |
